# Structural basis for phosphatidylcholine synthesis by bacterial phospholipid *N*-methyltransferases

**DOI:** 10.1101/2024.11.28.625837

**Authors:** Yasunori Watanabe, Hiroyuki Kumeta, Seiya Watanabe

## Abstract

In phosphatidylcholine (PC)-containing bacteria, PC is synthesized by phospholipid *N*-methyltransferases (Pmts) and plays an important role in the interactions between symbiotic and pathogenic bacteria and their eukaryotic host cells. Pmts catalyze the SAM dependent three methylation reactions of the head group of phosphatidylethanolamine (PE) to form PC through monomethyl PE and dimethyl PE. However, the precise molecular mechanisms underlying PC biosynthesis by PmtA remain largely unclear, owing to the lack of structural information. Here, we determined the crystal structures of *Agrobacterium tumefaciens* Pmt (AtPmtA) in complex with SAH or 5′-methylthioadenosine. Crystal structures and NMR analysis revealed the binding mode of AtPmtA to SAH in solution. Structure-based mutational analyses showed that a conserved tyrosine residue in the substrate-binding groove is involved in methylation. Furthermore, we showed that differences in substrate specificity among Pmt homologs were determined by whether the amino acid residues comprising the substrate-binding groove were isoleucine or phenylalanine. These findings provide a structural basis for understanding the mechanisms underlying Pmts-mediated PC biosynthesis.

## Introduction

Phosphatidylcholine (PC) is one of the most abundant phospholipids in the biological membranes of eukaryotic cells, and can be synthesized via two different pathways: the CDP-choline (or Kennedy) and *N*-methylation pathways (1). However, most bacterial membranes lack this phospholipid. For example, in the gram-negative model bacteria *Escherichia coli*, the cytoplasmic membrane consists mainly of three phospholipids: phosphatidylethanolamine (PE, 75%), phosphatidylglycerol (20%), and cardiolipin (5%) (2, 3). However, approximately 15% of all bacteria can produce PC; these include photosynthetic or symbiotic and pathogenic bacteria that interact with eukaryotic hosts (4–6). Bacterial PC functions primarily in interactions between symbiotic and pathogenic bacteria and their eukaryotic host cells (6–8). Moreover, mutants defective in PC synthesis have been reported in several bacterial species. Deletion of PC synthetic activity in the plant-transforming bacterium *Agrobacterium tumefaciens* prevents tumor formation in host cells because PC-deficient mutants lack the type IV secretion machinery required for T-DNA transfer to the host cells (9). In the microsymbiotic nitrogen-fixing bacterium *Bradyrhizobium diazoefficiens* (formerly *Bradyrhizobium japonicum*), a reduction in PC levels impairs efficient symbiosis with its soybean host (10).

In PC-forming bacteria, this phospholipid is mainly synthesized by the PC synthase or *N*-methylation pathways or both. PC synthase, an integral membrane protein, catalyzes the conversion of CDP-diacylglycerol and choline to PC and CMP in the PC synthase pathway (11, 12). In the *N*-methylation pathway, phospholipid *N*-methyltransferases (Pmts), which are cytosolic proteins, catalyze the methylation of the head group of phosphatidylethanolamine (PE) using SAM as a methyl group donor to form PC via monomethyl (MMPE) and dimethyl PE (DMPE) (13).

Pmts are encoded by several PC-forming bacteria and classified into two groups based on sequence similarity: the *Rhodobacter*-type (R-type) and *Sinorhizobium*-type (S-type) (5). *A. tumefaciens* has an S-type Pmt called PmtA, which was most intensively characterized by Narberhaus et al.. This PmtA catalyzes all three methylation reaction steps required for PC formation from PE via MMPE and DMPE (14, 15). Pmt homologs possess an N-terminal amphipathic helix that is involved in membrane binding and remodeling, along with a Rossmann fold domain containing a SAM-binding motif. (16–19). *B. diazoefficiens* possesses PmtA (BdPmtA) and three additional Pmts (BdPmtX1, BdPmtX3, and BdPmtX4). However, Pmts from *B. diazoefficiens* differ in the substrate specificity of the methylation reaction for PC synthesis. S-type BdPmtA predominantly catalyzes the methylation of PE to MMPE, whereas R-type BdPmtX1 uses MMPE as a substrate to form DMPE and PC (10, 20, 21). R-type PmtA from thermophilic bacteria, *Rubellimicrobium thermophilum*, catalyzes all three methylation reactions (22).

Recently, the crystal structure of the *R. thermophilum*-derived R-type PmtA with DMPE and SAH provided insights into substrate recognition, membrane binding, and catalytic mechanisms (23). However, the precise molecular mechanisms underlying PC biosynthesis and the substrate specificity of S-type PmtA remain largely unclear because of the lack of its structural information. In this study, we report the structural and functional analyses of *A. tumefaciens* PmtA (AtPmtA). We determined the crystal structures of AtPmtA lacking the N-terminal amphipathic helix in complex with SAH or 5’-methylthioadenosine at 1.96 Å and 2.04 Å resolutions, respectively. NMR analyses of apo and SAH-bound forms of AtPmtA showed the precise molecular mechanism of SAM recognition. Structure-based mutational analyses provided the structural insights into the differences in substrate specificity of PmtA between *A. tumefaciens* and *B. diazoefficiens*. Our findings provide a structural basis for the molecular mechanisms underlying PC biosynthesis and the differences in substrate specificity among S-type PmtA homologs.

## Results

### Crystal structure of AtPmtA

AtPmtA consists of 197 amino acids and has an N-terminal amphipathic helix (amino acid 5– 24) that is responsible for membrane binding (17). In addition, it has a C-terminal catalytic domain containing a highly conserved SAM-binding motif (E/DXGXGXG; amino acid 58–64) (24). To elucidate the structural mechanism underlying PC synthesis by AtPmtA, we first attempted to determine its crystal structure at full length. However, it failed to crystallize. Because the N-terminal amphipathic helix is involved in membrane binding and is predicted to be intrinsically disordered, six N-terminal truncated mutants were constructed: ΔN5, ΔN10, ΔN15, ΔN20, ΔN25, and ΔN30 lacking the N-terminal residues 5, 10, 15, 20, 25, and 30, respectively. Among these constructs, the solubility and expression levels of AtPmtAΔN20, AtPmtAΔN25, and AtPmtAΔN30 in *E. coli* cells were markedly improved (Fig. S1). Thus, we used the three constructs for crystallization trials. We successfully obtained well-diffracted crystals of AtPmtAΔN25 through co-crystallization with SAH. The crystal structure of AtPmtAΔN25 containing SAH was determined by the molecular replacement method using the structure predicted by AlphaFold 2 as the search model (25, 26). We subsequently refined it to a resolution of 1.96 Å (Table 1).

**Table 1.** Data collection, phasing and refinement statistics.

|  | AtPmtAΔN25-SAH | AtPmtAΔN25-MTA |
| --- | --- | --- |
| **Data collection** |  |  |
| Space group | $P2_1$ | $P2_1$ |
| Cell dimensions |  |  |
| $a, b, c$ (Å) | 69.94, 76.38, 87.19 | 69.61, 76.39, 87.23 |
| $\alpha, \beta, \gamma$ (°) | 90.0, 105.8, 90.0 | 90.0, 105.6, 90.0 |
| Data set |  |  |
| Wavelength (Å) | 1.00000 | 1.00000 |
| Resolution range (Å) | 50.0–1.95 (2.07–1.95) | 50.0–2.04 (2.16–2.04) |
| $R_{\text{merge}}$ | 0.348 (2.105) | 0.322 (2.055) |
| $R_{\text{meas}}$ | 0.387 (2.333) | 0.358 (2.275) |
| $I/\sigma I$ | 6.8 (1.0) | 7.1 (1.1) |
| Completeness (%) | 99.5 (97.9) | 99.3 (98.1) |
| Redundancy | 5.2 (5.3) | 5.3 (5.4) |
| $CC_{1/2}$ | 0.979 (0.454) | 0.984 (0.409) |
| **Refinement** |  |  |
| Resolution (Å) | 46.6–1.95 | 46.7–2.04 |
| $R / R_{\text{free}}$ | 0.187 / 0.227 | 0.170 / 0.209 |
| No. atoms |  |  |
| Protein | 5162 | 5145 |
| Ligand | 104 | 80 |
| Water | 620 | 499 |
| Sulfate ion | 5 | - |
| $B$-factors (Å<sup>2</sup>) | | |
| Protein | 28.6 | 31.0 |
| Ligand | 27.9 | 35.8 |
| Water | 36.7 | 56.8 |
| Sulfate ion | 40.6 | - |
| R.m.s. deviations |  |  |
| Bond lengths (Å) | 0.007 | 0.007 |
| Bond angles (°) | 0.8 | 0.8 |
| Ramachandran plot |  |  |
| Favored (%) | 98.0 | 97.4 |
| Allowed (%) | 2.0 | 2.6 |
| Outliers (%) | 0.0 | 0.0 |
| PDB accession code | 9KO3 | 9KO5 |

Values in parentheses represent the highest-resolution shell.

The crystallographic asymmetric unit contains four AtPmtAΔN25 molecules that have similar conformations, with a root-mean-square deviation (r.m.s.d.) of 0.18–0.30 Å for 135 Cα atoms. Three of the four AtPmtAΔN25 models lacked eight N-terminal residues (residues 26-33) because of an undefined electron density, which is consistent with the prediction that the N-terminal is intrinsically disordered. In addition, AtPmtAΔN25 consisted of a core Rossman-fold domain composed of a seven-stranded β-sheet (β1-β7) surrounded by five α-helices (α1-α5) (Fig. 1A). A query to the DALI server (27) revealed several SAM-dependent methyltransferases that were structurally similar to AtPmtAΔN25. The DALI search identified the following methyltransferases as structurally related: the tRNA:m^2^G6 methyltransferase TrmN (PDB code 3TMA; 2.4 Å r.m.s.d.; 15% identity) (28), RsmD-like methyltransferase (PDB code 3P9N; 2.8 Å r.m.s.d.; 18% identity) (29), tetrahydroprotoberberine *N*-methyltransferase (PDB code 6P3O; 2.7 Å r.m.s.d.; 12% identity) (30), and phosphoethanolamine methyltransferase (PDB code 3UJ7; 2.7 Å r.m.s.d.; 17% identity) (31) from *Thermus thermophilus*, *Mycobacterium tuberculosis*, *Glaucium flavum*, and *Plasmodium falciparum*, respectively. All four AtPmtAΔN25 molecules in the asymmetric unit contained SAH molecules with clear electron densities. Similar to other SAM-dependent methyltransferases, SAH bound close to the conserved Gly-rich loop region between β1 and α2 (Fig. 1B). Furthermore, we successfully co-crystallized AtPmtAΔN25 with SAM under similar crystallization conditions as the SAH-bound form of AtPmtAΔN25. The structure was determined at 2.04 Å resolution through molecular replacement using the structure of SAH-bound AtPmtAΔN25 as the search model (Table 1). Only partial electron density of the SAM molecule was observed in the crystal structure (Fig. 1C). Due to SAM instability, it undergoes cleavage to 5′-methylthioadenosine (MTA) and homoserine lactone (32). Therefore, we assigned the electron density to MTA.

**Figure 1.**
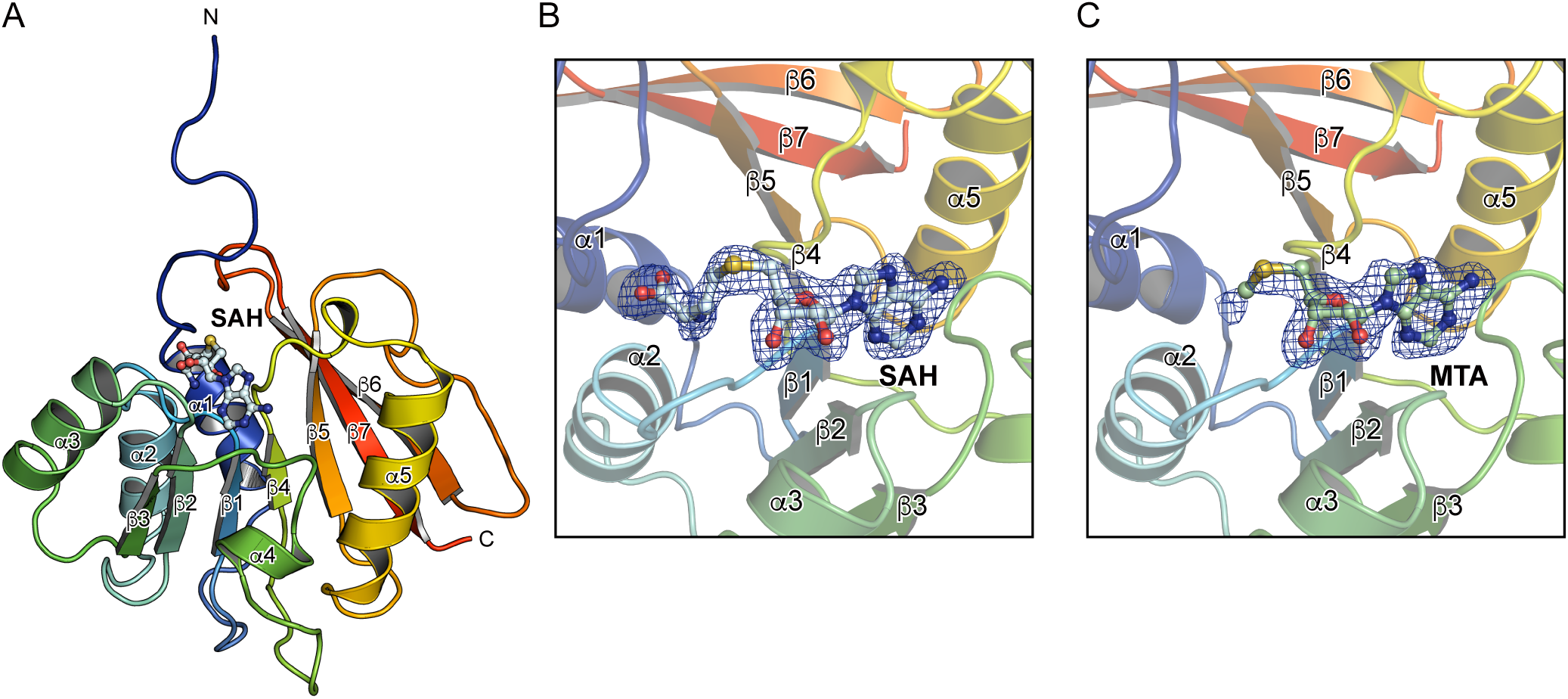
AtPmtAΔN25 structure. (A) A ribbon diagram of the SAH-bound AtPmtAΔN25 structure, colored blue to red from the N to the C terminus. Secondary structural elements are labeled. The bound SAH molecule is shown in stick and ball representation. (B-C) Electron density maps of SAH (B) and MTA (C). The simulated annealing *mF*o-*DF*c difference Fourier maps were calculated by omitting the SAH and MTA molecules and are shown as blue meshes contoured at 3.0σ.

### NMR analysis of the interaction of AtPmtA with SAH

We performed NMR studies on AtPmtAΔN25 to elucidate the mechanism underlying SAH recognition by AtPmtA in solution. The ^1^H-^15^N HSQC spectra of the apo- and SAH-bound forms of AtPmtAΔN25 with backbone resonance assignments are shown in Fig. S2. Both forms of AtPmtAΔN25 exhibited broad chemical shift dispersion in the ^1^H-^15^N HSQC spectra, suggesting that they both form a well-folded domain. However, resonances corresponding to 21 of the 157 non-proline residues in the apo-form of AtPmtAΔN25 were missing from the ^1^H- ^15^N HSQC spectrum. The missing assignments may be due to the fast exchange of amide protons with the solvent or local exchange broadening. In contrast, resonances corresponding to 14 of the 21 AtPmtAΔN25 residues appeared in the ^1^H-^15^N HSQC spectrum upon binding with SAH. Eight of the 14 residues for which NMR signals were observed upon SAH binding clustered around the SAM-binding motif (E/DXGXGXG; amino acid 58–64) (Fig. 2A). Furthermore, some NMR signals showed considerable chemical shift changes upon SAH binding (Fig. 2A). Residues whose resonances appeared upon binding to SAH and those exhibiting considerable chemical shift chenges were mapped to the crystal structure of AtPmtAΔN25-SAH (Fig. 2B). These residues were clustered at the binding interface with SAH. These observations suggest SAH-induced conformational changes in AtPmtAΔN25 to a stable conformation, consistent with the requirement of SAH or SAM for the crystallization of AtPmtAΔN25.

**Figure 2.**
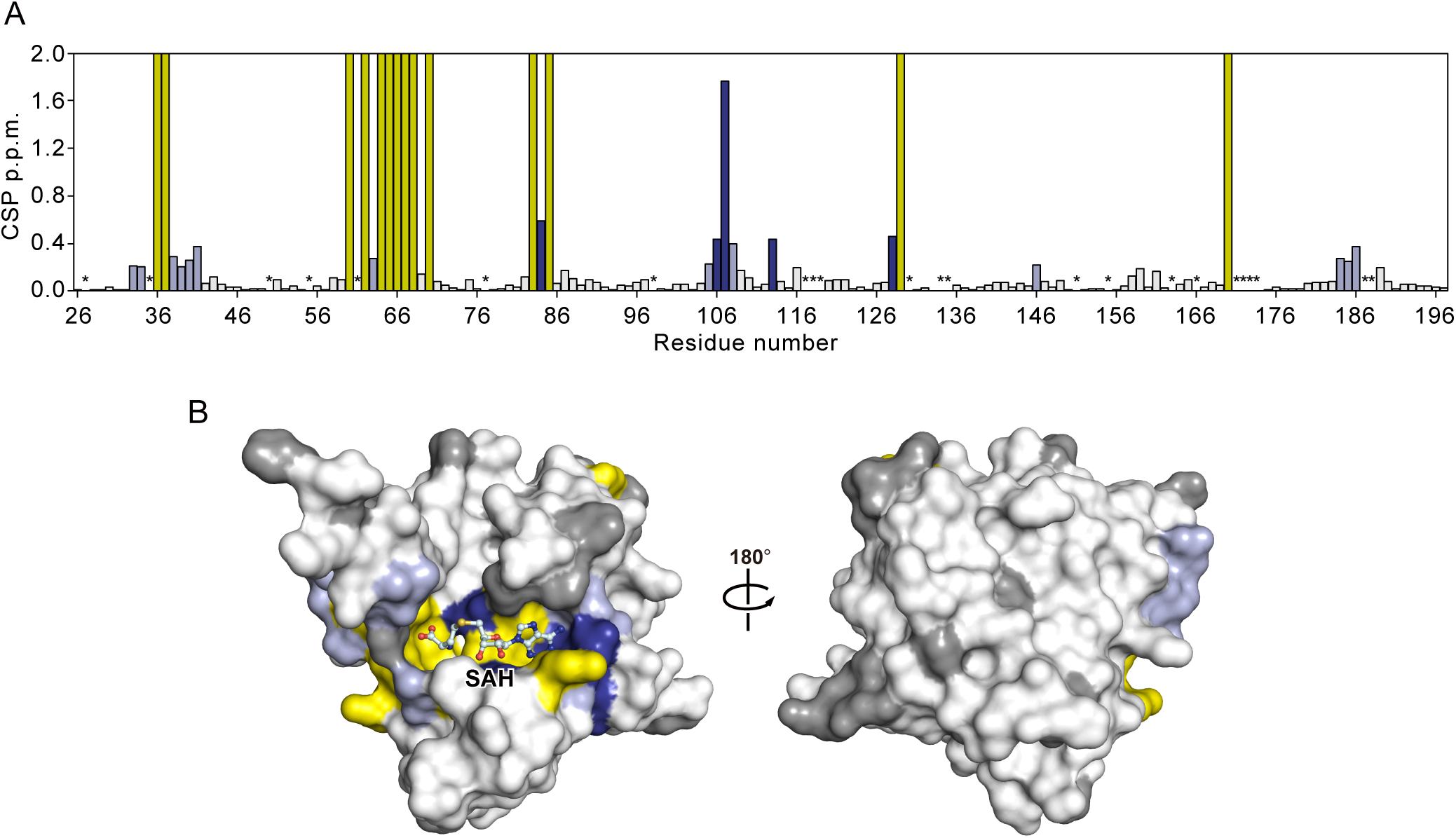
NMR analysis of SAH binding of AtPmtA. (A) Chemical shift perturbation (CSP) values of amide moieties of AtPmtAΔN25 upon binding to SAH plotted for each residue on AtPmtAΔN25. The residues with CSPs higher than the threshold values of 0.2 and 0.4 ppm are colored light and dark blue, respectively. The residues whose signals appeared upon binding to SAH are colored yellow. Proline and unassigned residues are indicated by asterisks. (B) The residues with significant CSPs upon binding to SAH are mapped on the surface of the crystal structure of SAH-bound AtPmtAΔN25 using the same color representations as in (A). The proline and unassigned residues are colored grey. SAH is shown in stick form.

### Structural basis of the AtPmtAΔN25-SAH interaction

The detailed interaction between AtPmtAΔN25 and SAH is shown in Figure 3A. The adenine group of SAH is well-recognized by Glu84, Tyr85, Asp106, Ala107, Phe108, and Val129 in the crystal structure of the AtPmtAΔN25-SAH complex. In addition, it is within a distance that allows the formation of hydrogen bonds with the side chain of Asp106 and the main chain nitrogen atoms of Glu84 and Ala107. The ribose and homocysteine groups of SAH were located near Thr36, Gly60, Gly62, Val65, Ile66, Glu84, and Ala128. Moreover, the 2’ and 3’ hydroxyls of the ribose group were within a distance that allows the formation of hydrogen bonds with the side chain of Glu84. The main chain of Gly60 and side chain of Thr36 can form hydrogen bonds with the homocysteine group. To further investigate the cofactor recognition mechanism, we substituted Thr36, Glu84, and Asp106, which can form hydrogen bonds with the homocysteine, ribose, and adenine groups, respectively, and with alanine residues (T36A, E84A, and D106A). Ala107, which is located near the adenine group, exhibited the largest chemical shift upon SAH binding (Fig. 2A). Therefore, we constructed a mutant in which Ala107 was substituted with a phenylalanine residue (A107F). The enzyme activity of AtPmtA mutants was assessed by expressing them in *E. coli* cells without MMPE, DMPE, and PC. Expression patterns of AtPmtA mutants are shown in Figure 3B. Additionally, total phospholipids in AtPmtA mutants-expressing cells are shown in Figure 3C. PE was primarily detected in *E. coli* cell membranes harboring the empty vector, but the methylated PE derivatives, MMPE, DMPE, and PC, were not detected. In contrast, the membranes of *E. coli* cells expressing wild-type AtPmtA contained PE, MMPE, DMPE, and PC. However, only trace amounts of DMPE were present, indicating that wild-type AtPmtA catalyzed all three methylations from PE to PC. *E. coli* cell membranes expressing the AtPmtA T36A mutant also contained PE, MMPE, DMPE, and PC, although the amount of PC in *E. coli* cells expressing the AtPmtA T36A mutant was slightly lower than that in cells expressing wild-type AtPmtA. This suggests that the T36A mutation had little effect on the catalytic activity of AtPmtA. PE methyltransferase activity was almost completely lost in the E84A and D106A mutations and dramatically reduced in the A107F mutation. This suggests that Glu84, Asp106, and Ala107 are important for PE methyltransferase activity.

**Figure 3.**
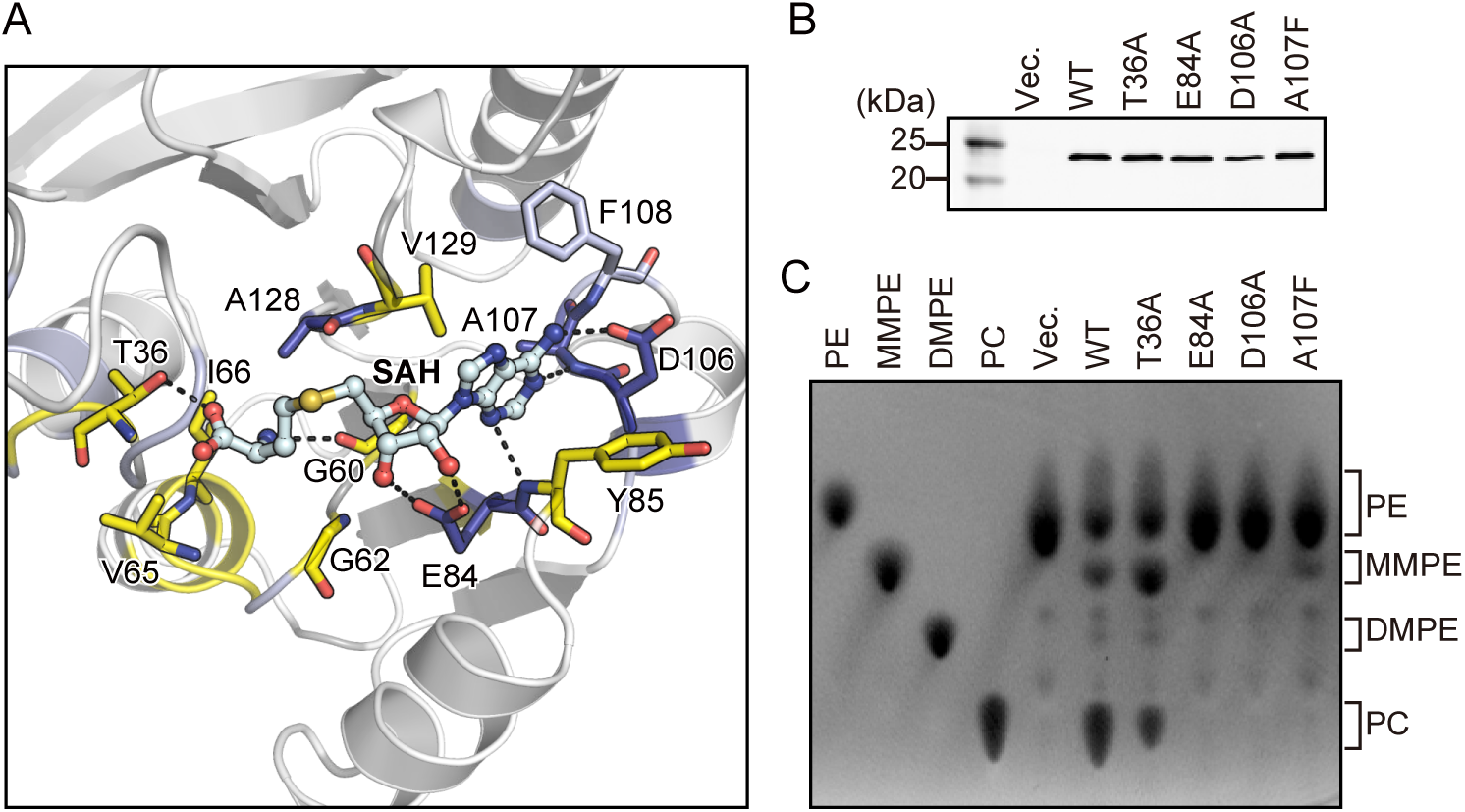
Residues responsible for SAH binding. (A) A magnified view showing the detailed interaction around SAH. Residues responsible for SAH recognition are shown as stick models using the same color representations as that in Fig. 2C. Broken lines represent possible hydrogen bonds. (B) Lysates from *E. coli* cells expressing N-terminal His_6_-tagged AtPmtA mutants were subjected to SDS-PAGE, followed by immunoblotting with anti-6x histidine antibody. (C) Total phospholipids were extracted from *E. coli* cells expressing N-terminal His_6_-tagged AtPmtA mutants and separated using TLC, followed by detection using 2’,7’-dichlorofluorescein. 18:1-18:1 PE, 18:1-18:1 MMPE, 18:1- 18:1 DMPE and 18:1-18:1 PC were used as TLC markers.

### Putative substrate phospholipid binding site

In order to investigate the substrate recognition mechanism, we tried to determine the ternary complex crystal structure of AtPmtA with SAH and PE or PC. However, we could not obtain crystals of the ternary complex by soaking or co-crystallization. Therefore, we performed a docking simulation using the AutoDock Vina suite (33) to model the ternary complex of AtPmtAΔN25-SAH and phosphocholine (pCho), a PC head group. pCho was docked to the putative substrate-binding site near SAH in the crystal structure of the AtPmtAΔN25-SAH complex (Fig. 4A). In the docked model, the nitrogen atom of pCho is close to the sulfur atom of SAH, and the distance between two atoms is 3.9 Å (Fig. 4B). This suggests that the docking model is reasonable. pCho is located near the main chains of Ala32, Ile33, Val34, Pro35, Thr36, Ala128, and Ile186 and the side chains of Thr36, Thr63, Phe89, Ala128, Ile159, and Tyr161. Additionally, the trimethylamine moiety is accommodated in the grooves formed by Ala128, Ile159, and Tyr161 (Fig. 4C).

**Figure 4.**
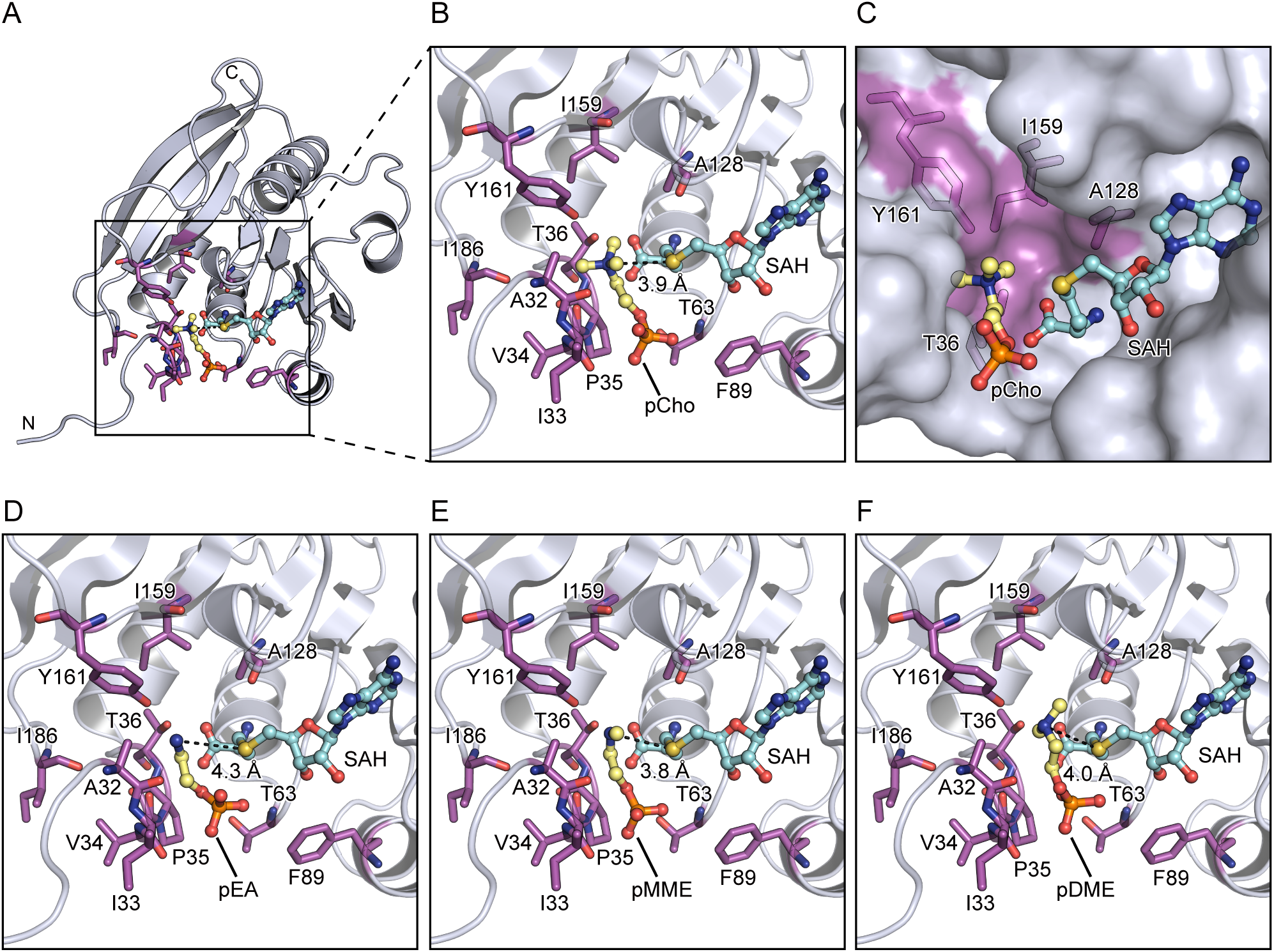
Substrate phospholipid head group docking models. (A) A docking model of the AtPmtAΔN25-SAH complex with pCho. pCho and SAH are shown in stick and ball representation. (B) A magnified view around pCho. Residues in close proximity to pCho are shown in stick form and labeled in magenta. The distance between the nitrogen and sulfur atoms of pCho and SAH, respectively, is indicated by a broken line. (C) A surface model of the docking site showing the pCho accommodated to the groove of AtPmtAΔN25. Thr36, Ala128, Ile159, and Tyr161, which are forming the groove, are shown in stick form and labeled in magenta. (D–F) Magnified views around the substrate phospholipid head group molecules in the docking models of the AtPmtAΔN25-SAH complex with pEA (D), pMMEA (E), and pDMEA (F). Residues indicated in (B) are shown in stick form and labeled in magenta. The distances between the nitrogen atom of the substrate phospholipid head group molecules and the sulfur atom of SAH are indicated by a broken line.

We also performed a docking simulation to model the ternary complex of AtPmtAΔN25-SAH and the following phospholipid head group molecules: phosphatidylethanolamine (pEA), phosphomonomethylethanolamine (pMMEA), and phosphodimethylethanolamine (pDMEA) (Fig. 4D–F). In the docking models, the nitrogen atoms of pEA, pMMEA, and pDMEA were close to the sulfur atom of SAH, and the distances between two atoms were 4.3 Å, 3.8 Å, and 4.0 Å, respectively. Similar to pCho, the amine moieties of pEA, pMMEA, and pDMEA were accommodated in the grooves formed by Ala128, Ile159, and Tyr161. These results suggest that AtPmtA catalyzes a three-step methylation reaction at a common substrate-binding site.

### Substrate phospholipid recognition mechanism

AtPmtA catalyzes a three-step methylation reaction that produces PC from PE via MMPE and DMPE. In contrast, BdPmtA predominantly catalyzes the methylation of PE to MMPE with little subsequent reaction with DMPE and PC (20). Therefore, we compared the crystal structure of the AtPmtAΔN25-SAH complex and the AlphaFold 2-predicted structure of BdPmtA to investigate the substrate specificity of AtPmtA and BdPmtA. The predicted BdPmtA structure is similar to the crystal structure of the AtPmtAΔN25-SAH complex (1.2 Å r.m.s.d. for 148 Cα atoms; 32% identity) (Fig. 5A). Among the residues that may be involved in the recognition of the amine moiety of substrate phospholipids, Tyr161 was conserved (Tyr163 in BdPmtA). However, Ala128 and Ile159 were substituted with glycine and phenylalanine, respectively (G130 and F161 in BdPmtA) (Fig. 5B and C). We substituted tyrosine residues with alanine or phenylalanine (Y161A and Y161F in AtPmtA, or Y163A and Y163F in BdPmtA) and assessed the enzyme activity of AtPmtA or BdPmtA mutants by expressing them in *E. coli* cells. The expression patterns of these mutants in *E. coli* cells are shown in Figure 5D. TLC analysis of phospholipids from *E. coli* cells expressing AtPmtA mutants showed that methylation activity was severely reduced in the Y161A mutant and significantly reduced in the Y161F mutant (Fig. 5E). The expression of wild-type BdPmtA mainly led to the production of MMPE and slight production of DMPE. Similar to AtPmtA, the methylation activity of BdPmtA was almost completely lost in the Y163A mutant and significantly reduced in the Y163F mutant. These results suggest that the conserved tyrosine residues in the groove (Tyr161 in AtPmtA and Tyr163 in BdPmtA) are important for PE methylation activity.

**Figure 5.**
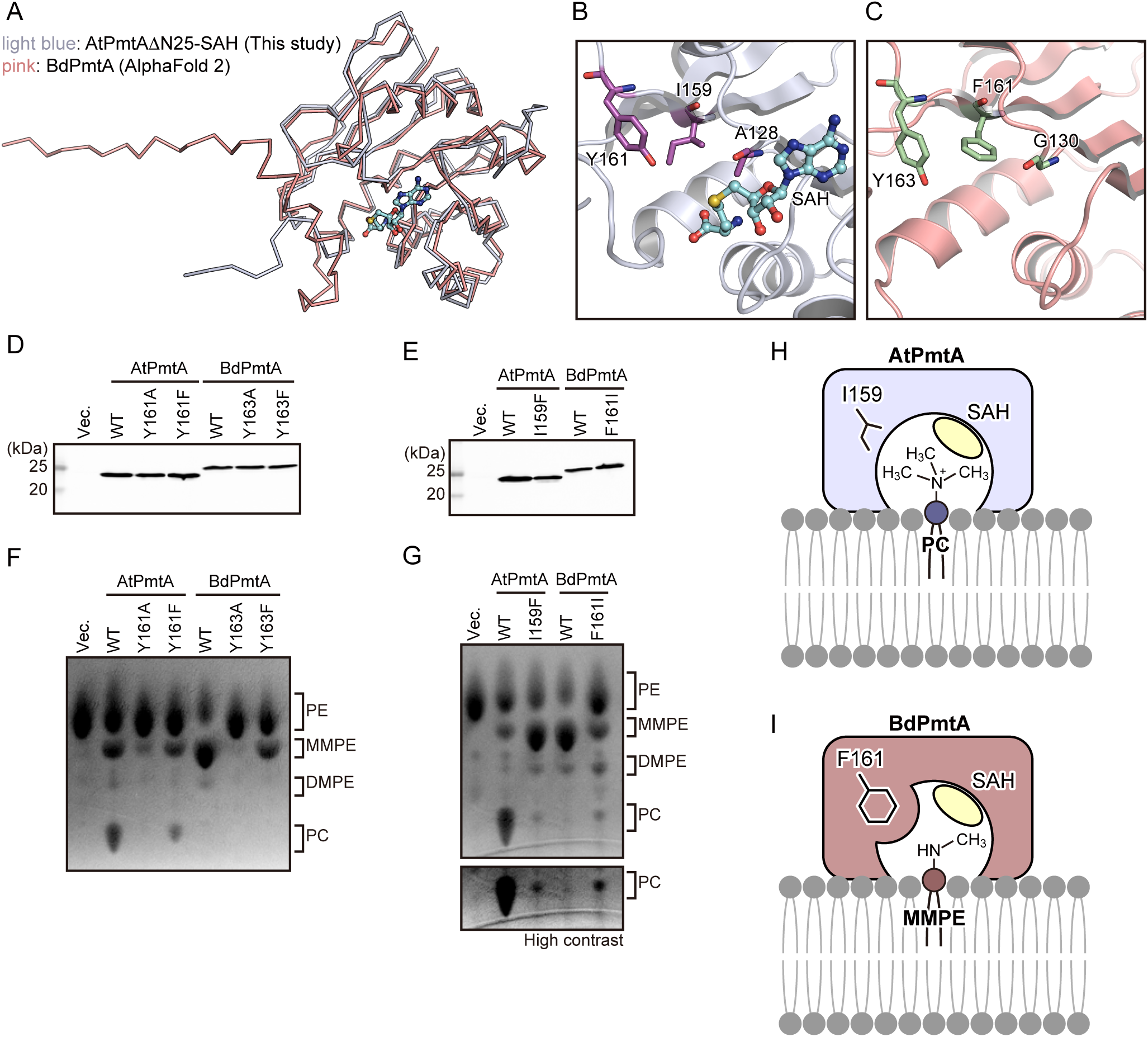
Substrate specificities of AtPmtA and BdPmtA. (A) Superposition of the crystal structure of the AtPmtAΔN25-SAH complex and the AlphaFold 2-predicted structure of BdPmtA. AtPmtAΔN25 and BdPmtA are colored in light blue and pink, respectively. SAH bound to AtPmtAΔN25 in shown in stick and ball representation. (B-C) Magnified views of the putative substrate binding sites of AtPmtA (B) and BdPmtA (C). Residues suggested to be involved in the recognition of the amine moiety of substrate phospholipids are shown in stick forms and labeled in magenta. (D) Lysates from *E. coli* cells expressing N-terminal His_6_-tagged AtPmtA mutants were subjected to SDS-PAGE, followed by immunoblotting with anti-6x histidine antibody. (E) Total phospholipids were extracted from *E. coli* cells expressing N-terminal His_6_-tagged AtPmtA mutants and separated using TLC, followed by detection using 2’,7’-dichlorofluorescein. (F) Lysates from *E. coli* cells expressing N-terminal His_6_-tagged BdPmtA mutants were subjected to SDS-PAGE, followed by immunoblotting with anti-6x histidine antibody. (G) Total phospholipids were extracted from *E. coli* cells expressing N-terminal His_6_-tagged BdPmtA mutants and separated using TLC, followed by detection using 2’,7’-dichlorofluorescein. A part of a TLC plate showing PC is shown with high contrast in the lower panel. (H–I) Putative phospholipid binding models of AtPmtA (H) and BdPmtA (I). The substrate phospholipid-binding groove containing Ile159 of AtPmtA can accommodate the head group of PC in the membrane. In contrast, the substrate-binding groove containing Phe161 of BdPmtA cannot accommodate the head group of PC, owing to steric inhibition, but can accommodate the head group of MMPE.

To evaluate the roles of Ile159 in AtPmtA and Phe161 in BdPmtA in substrate specificity, we substituted these residues with phenylalanine and isoleucine, respectively (I159F in AtPmtA and F161I in BdPmtA). The expression levels of these mutants in *E. coli* cells are shown in Figure 5F. TLC analysis of *E. coli*-derived phospholipids showed that expression of the AtPmtA I159F mutant led to the accumulation of MMPE and reduction of PC compared to that of wild-type AtPmtA (Fig. 5G). However, expression of the BdPmtA F161I mutant led to an increase in DMPE and PC and a reduction in MMPE compared to wild-type BdPmtA. These results indicate that Ile159 in AtPmtA and Phe161 in BdPmtA are responsible for substrate specificity.

## Discussion

The phospholipid *N*-methylation pathway, which is catalyzed by SAM-dependent phospholipid *N*-methyltransferases, is a major PC biosynthesis pathway in PC-containing bacteria. In this study, we primarily investigated the S-type AtPmtA and determined its crystal structure using SAH or MTA. We also performed NMR analysis of AtPmtA in the apo- and SAH-bound forms and structure-based mutational analyses using *E. coli* cells expressing PmtA mutants. Although we used AtPmtAΔN25, which lacks the N-terminal amphipathic helix involved in membrane binding, for X-ray crystallography and NMR analyses, structure-based functional analyses have provided structural insights into the molecular mechanisms underlying PC biosynthesis, substrate recognition, and specificity of S-type PmtA.

Our substrate-docking models indicated that the hydroxyl group of Tyr161 in AtPmtA is close to the amine moiety of the substrate phospholipids (Fig 4). Salsabila and Kim (23) determined the crystal structure of *R. thermophilum*-derived R-type PmtA in complex with SAH and DMPE and revealed that the tyrosine residue near the substrate phospholipids was involved in the activation of the amine through deprotonation. SAM-dependent phosphoethanolamine methyltransferases (PMTs), which catalyze the methylation of phosphoethanolamine to phosphocholine for membrane biogenesis in plants, nematodes, and *Plasmodium* apicomplexan parasites, also require a hydroxyl group of tyrosine residues near the amine of the substrate for methylation (31, 34, 35). Our mutational analyses showed that Tyr161 in AtPmtA and Tyr163 in BdPmtA played important roles in methylation (Fig. 5E), suggesting that conserved tyrosine residues in S-type PmtA are also involved in the activation of the amine moiety of substrate phospholipids. However, mutations in these tyrosine residues did not completely abolish methylation. Similar to PMTs from *P. falciparum* and *Haemonchus contortus* (31, 34), water molecules in the active sites may be involved in the deprotonation of the amine of the substrate.

We also found that Ile159 in AtPmtA and Phe161 in BdPmtA were involved in substrate specificity (Fig. 5G). AtPmtA and BdPmtA form substrate-binding grooves comprising isoleucine (Ile159) and phenylalanine (Phe161) residues, respectively. The substrate-binding groove of AtPmtA was wide enough to accommodate the head groups of DMPE and PC (Fig. 5H). In contrast, the substrate-binding groove of BdPmtA was narrow; therefore, it did not accommodate the head groups of DMPE and PC (Fig. 5I). Functional analyses of the substrate specificity of several S-type PmtA homologs have been reported. PmtA from *Sinorhizobium meliloti* (SmPmtA) catalyzes all three methylation reactions from PE to PC (36). BdPmtX3 catalyzes two methylation reactions from PE to DMPE, whereas BdPmtX4 and PmtA from *Xanthomonas campestris* (XcPmtA) catalyze predominantly the initial methylation from PE to MMPE (20, 37). SmPmtA and BdPmtX3 possess isoleucine residues equivalent to Ile159 in AtPmtA, whereas BdPmtX4 and XcPmtA possess phenylalanine residues equivalent to Phe161 in BdPmtA. These findings indicate that the substrate specificity of S-type PmtA is determined by the size of the substrate-binding groove, which is defined by whether the residues in the groove are isoleucine or phenylalanine.

In yeast, the integral membrane enzymes Cho2 and Opi3 catalyze three SAM-dependent methylation reactions of PE to synthesize PC. Cho2 catalyzes the first methylation of PE to synthesize MMPE, whereas Opi3 catalyzes the second and third methylations to synthesize PC via DMPE from MMPE (38, 39). In mammals, PEMT, which is a PE *N*-methyltransferase homologous to Opi3, catalyzes all three SAM-dependent methylations of PE to PC (40). Eukaryotic PE *N*-methyltransferases such as bacterial Pmts may also have substrate specificity, which is determined by the size of the substrate-binding region. Therefore, future structural studies on eukaryotic PE *N*-methyltransferases provide the structural basis of their substrate specificity.

In this study, we determined the crystal structure of AtPmtA through the molecular replacement method using the AlphaFold 2-predicted structure as the search model. The predicted structure of AtPmtA was markedly similar to the crystal structure of AtPmtA in the Rossmann fold domain (r.m.s.d. of 0.426 Å for 148 Cα atoms) but different in the N-terminal (Fig. S4). Ile33 in the predicted structure was too close to the SAM-binding region, whereas it was further away from the SAM-binding region in the crystal structure. This suggests that the actual orientation of the N-terminal amphipathic helix was different from that in the predicted structure. The N-terminal amphipathic helix of AtPmtA is required for PE methylation, membrane binding, and remodeling (Fig. S1) (17–19). Salsabila and Kim (23) showed that the membrane binding of R-type PmtA from *R. thermophilum* required not only the N-terminal helix, but also other helices, which are conserved in R-type PmtA homologs. Except for the N-terminal helix, the helices involved in membrane binding are not structurally conserved in AtPmtA. This suggests that the N-terminal helix of AtPmtA is the only membrane-binding region, and that membrane-binding mechanisms differ between AtPmtA and R-type PmtA. Thus, the full-length structure of AtPmtA, including the N-terminal helix, requires further study to elucidate the molecular mechanism underlying PC biosynthesis at the membrane-water interface.

In conclusion, the crystal structure of AtPmtA provides not only a structural basis for the molecular mechanism underlying PE methylation and the substrate specificity of S-type PmtA, it also paves the way for future studies on biogenesis of cell membrane and endomembrane.

## Experimental procedures

### Construction of expression plasmids

The AtPmtA DNA fragment was amplified from the genome of *A. tumefaciens* NBRC 15193 using PCR and cloned into the pETDuet-1 vector to construct *E. coli* expression plasmids for full-length AtPmtA. To construct *E. coli* expression plasmids for AtPmtAΔN25 (lacking a.a. 1–25) with a cleavable N-terminal His_6_-tag, the DNA fragment for AtPmtAΔN25 was amplified via PCR using pETDuet-1-AtPmtA as the template and cloned into the pETDuet-1 vector. The human rhinovirus (HRV) 3C protease recognition site was then inserted between the His_6_-tag and AtPmtAΔN25 in this plasmid using inverse PCR. The BdPmtA DNA fragment was amplified from the genome of *B. diazoefficiens* and cloned into the pETDuet-1 vector to construct *E. coli* expression plasmids for BdPmtA. Mutations in the described amino acid substitutions were introduced using PCR-based site-directed mutagenesis. All constructs were sequenced to confirm their identities.

### Protein expression and purification

AtPmtAΔN25 was expressed in *E. coli* C43 (DE3) cells (Lucigen), cultured in LB medium supplemented with 100 μg/mL ampicillin at 37 °C. When OD_600_ reached approximately 0.8, IPTG was added to a final concentration of 0.1 mM and cultured at 25 °C for 18 h to induce protein expression. Next, the cells were harvested, resuspended in lysis buffer (50 mM Tris-HCl [pH 8.0], 500 mM NaCl, and 20 mM imidazole), disrupted via sonication, and centrifuged at 20,000 *×g* for 40 min to pellet the insoluble debris. The supernatant was loaded onto a Ni-NTA agarose column (QIAGEN) equilibrated with lysis buffer. The column was washed with wash buffer (50 mM Tris-HCl [pH 8.0], 500 mM NaCl, and 20 mM imidazole), and His-tagged AtPmtAΔN25 was eluted using elution buffer (50 mM Tris-HCl [pH 8.0], 100 mM NaCl, and 250 mM imidazole). The eluate was treated overnight with HRV 3C protease at 4 °C to cleave the His_6_-tag. The protein was purified using a HiTrap SP HP cation-exchange column (GE Healthcare) with a linear gradient of 0–1000 mM NaCl in 50 mM sodium phosphate (pH 7.0). It was further purified through size exclusion chromatography using a Superdex 200 Increase column (GE Healthcare) with an elution buffer of 20 mM Tris-HCl (pH 8.0) and 150 mM NaCl. For NMR analyses, uniformly labeled AtPmtAΔN25 was expressed and purified as described above, except that an M9 medium containing ^15^N-ammonium chloride and [D-^13^C] glucose was used. The NMR samples were prepared in 20 mM HEPES-NaOH (pH 7.0) and 150 mM NaCl.

### Crystallization and X-ray crystallography

A crystallization trial was performed at 20 °C using the sitting drop vapor diffusion method. A purified AtPmtAΔN25 solution was mixed with SAH or SAM at a 10:1 ligand-to-protein molar ratio to prepare the complex between AtPmtAΔN25 and SAH or SAM. Drops (0.5 μL) of the mixture (approximately 27 mg/mL AtPmtAΔN25 in 20 mM Tris-HCl [pH 8.0] and 150 mM NaCl) were mixed with a reservoir solution (0.2 M lithium sulfate, 0.1 M Tris-HCl [pH 8.0], and 25% polyethylene glycol 3350) and equilibrated against 70 μL of the same reservoir solution using vapor diffusion. The crystals were soaked in a reservoir solution supplemented with 15% ethylene glycol, flash-cooled, and stored in a stream of nitrogen gas at 100 K during data collection. X-ray diffraction data were collected at the SPring-8 beamline BL32XU with a 10 *×* 15 μm (width *×* height) microbeam using the helical data collection method. The diffraction data were collected using an automated data collection system ZOO (41). Data processing was performed using the KAMO (42) and XDS (43) software. Structures were determined through molecular replacement using PHASER (44), for which the structure predicted by AlphaFold 2 (25, 26) was used as a search model. Further model building was performed manually using COOT (45), and crystallographic refinement was performed using PHENIX (46). MolProbity (47) was used to assess the quality and geometry of structural models. Detailed data collection and processing statistics are shown in Table1.

### NMR spectroscopy

^13^C/^15^N-labeled AtPmtAΔN25 (0.3 mM) was prepared in 20 mM HEPES-NaOH (pH 7.0), 150 mM NaCl, 0 or 3 mM SAH, and 10% d6-DMSO. NMR experiments were performed using a Bruker AVANCE NEO 800 MHz spectrometer equipped with a CPTCI probe, a Bruker AVANCE III HD 600 MHz spectrometer equipped with a TBI probe, and an Agilent Unity INOVA 600 MHz NMR spectrometer equipped with a TR5 probe. The experiments were performed at 10 °C for the apo-form of AtPmtAΔN25 and at 25 °C for SAH-bound AtPmtAΔN25. The spectra were processed using TopSpin (Bruker Co. Ltd.) and NMRPipe (48) and analyzed using Sparky software (Goddard and Kneller, Sparky 3, https://www.cgl.ucsf.edu/home/sparky/). Backbone assignments of apo- and SAH-bound AtPmtAΔN25 were obtained from the two-dimensional ^1^H-^15^N HSQC and three-dimensional HN(CO)CA, HNCA, CBCA(CO)NH, HNCACB, and ^15^N-NOESY (mixing time 150 ms) spectra. All three-dimensional spectra except for ^15^N-NOESY were recorded using the 25% non-uniform sampling (NUS) method. NUS spectra were reconstructed with a compressed sensing algorithm using qMDD (49) in the TopSpin program.

NMR titration experiments for SAH were performed at 10 °C. ^1^H-^15^N HSQC was measured at 0, 0.33, 0.67, 1, 2, 3, 5, and 10 molar equivalents of SAH conditions. Chemical shift changes in the amide moiety between SAH-free (0 eq.) and -bound (10 eq.) AtPmtAΔN25 were normalized using the following equation:

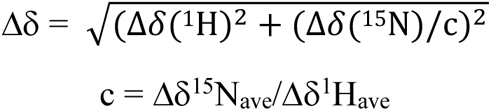

where Δδ^15^N_ave_ and Δδ^1^H_ave_ were average of Δδ^15^N and Δδ^1^H, respectively.

### In-silico docking

AutoDock Vina (33) was used as the docking tool to generate a putative AtPmtAΔN25-SAH complex bound to phosphocholine. The input files were prepared using AutoDock Tools. The AtPmtAΔN25-SAH complex structure was set as rigid during docking, and a 8 *×* 8 *×* 8 Å grid box was placed into the substrate-binding site. All AutoDock Vina parameters were maintained at their default values.

### Separation of soluble and membrane fractions

Separation of the soluble and membrane fractions from *E. coli* lysates was performed as previously described, with several modifications (50). *E. coli* C43(DE3) cells carrying the expression plasmids of AtPmtA or BdPmtA mutants were cultured in 50 mL LB medium at 37 °C. When the OD_600_ reached approximately 0.5, IPTG was added to a final concentration of 100 μM and the cultures were grown at 25 °C for 18 h to induce protein expression. Next, the cells were harvested, resuspended in lysis buffer (20 mM Tris-HCl [pH 8.0] and 150 mM NaCl), disrupted through sonication, and centrifuged at 5,000 *×g* for 10 min to pellet the insoluble debris. The samples were ultracentrifuged at 120,000 *×g* for 1 h. The supernatant and pellet fractions were defined as soluble and membrane fractions, respectively. They were then analyzed using SDS-PAGE and immunoblotting.

### TLC analysis of phospholipids

The membrane fractions from 50 mL *E. coli* cultures were suspended in 500 μL of lysis buffer using sonication to measure the PE *N*-methylation activity of PmtA mutants. One hundred microliters of the membrane fraction suspension was mixed with 900 μL chloroform/methanol (2:1, v/v) and vortexed for 15 min. Two hundred microliters of water was added to the samples, which were then vortexed for 10 min. The organic phase was separated via centrifugation at 1,000 ×*g* for 2 min, collected, and dried under N_2_ gas. The resulting lipid films were dissolved in chloroform (80 μL). Twenty-microliter aliquots of the samples and 20 μg each of 18:1-18:1 PE, 18:1-18:1 MMPE, 18:1-18:1 DMPE, and 18:1-18:1 PC were loaded and analyzed via TLC using *n*-propanol/propionic acid/chloroform/H_2_O (3:2:2:1). The TLC plate (Merck Millipore) was then dipped in ethanol containing 0.0002% 2’,7’-dichlorofluorescein for phospholipid detection. Fluorescence signals were detected using Typhoon FLA 9500 (GE Healthcare). Phospholipids were purchased from Avanti Polar Lipids, Inc.

### Immunoblotting

Mouse monoclonal anti-6x histidine antibody (9C11) was purchased from FUJIFILM Wako. The antibody was used in 1:2000 dilution for immunoblotting. Proteins were visualized with a fluorophore-conjugated secondary antibody, Goat anti-Mouse IgG (H + L) Cross-Absorbed Secondary Antibody, Cyanine5 (A10524; Life Technologies) used in 1:2000 dilution. The signals were analyzed using Typhoon FLA 9500 (GE Healthcare).

### Amino acid sequence alignment

The amino acid sequences of AtPmtA and its homologs were analyzed using Clustal Omega (51) and ESPript 3.0 (52).

## Data availability

The data used in this study are available upon request from the corresponding author. The atomic coordinates and structure factor files were deposited in the Protein Data Bank under the accession codes 9KO3 (AtPmtAΔN25 with SAH) and 9KO5 (AtPmtAΔN25 with MTA). The assignments of the backbone resonances were deposited in the Biological Magnetic Resonance Bank under the accession numbers 52730 (apo-form of AtPmtAΔN25) and 52731 (SAH-bound form of AtPmtAΔN25).

## Supporting information

This article contains supporting information.

## Acknowledgements

We thank Dr. Yasushi Tamura for his valuable advice and discussions. Synchrotron radiation experiments were performed at beamline BL32XU at SPring-8, Japan, with approval from the Japan Synchrotron Radiation Research Institute (Proposal Numbers 2020A2572, 2021A2762, and 2021B2762). NMR experiments were performed at the Hokkaido University NMR Platform, supported by the Ministry of Education, Culture, Sports, Science, and Technology (MEXT), Japan.

## Author contributions

Y.W. conceived the project and performed protein preparation, crystallographic studies and mutational analyses. H.K. performed NMR studies. S.W. contributed to the preparation of the expression plasmids. Y.W. wrote the manuscript. All authors reviewed and commented on the manuscript.

## Funding and additional information

This work was supported by the Japan Society for the Promotion of Science KAKENHI (Grant Numbers 22K06096 and 20K15734 to Y.W.), Sumitomo Foundation (to Y.W.), and Institute for Fermentation, Osaka (to Y.W.).

## Abbreviations and nomenclature

Pmt: phospholipid *N*-methyltransferase
PC: phosphatidylcholine
PE: phosphatidylethanolamine
AtPmtA: *Agrobacterium tumefaciens* Pmt
MMPE: monomethyl
DMPE: dimethyl PE
MTA: 5**′**-methylthioadenosine
pCho: phosphocholine
pEA: phosphatidylethanolamine
pMMEA: phosphomonomethylethanolamine
pDMEA: phosphodimethylethanolamine
PMTs: phosphoethanolamine methyltransferases
SmPmtA: PmtA from *Sinorhizobium meliloti*
XcPmtA: PmtA from *Xanthomonas campestris*
BdPmtA: PmtA from *B. diazoefficiens*
HRV: human rhinovirus
NUS: non-uniform sampling;
CSP: chemical shift perturbation.

## References

1. Cockcroft, S. (2021) Mammalian lipids: structure, synthesis and function Essays Biochem 65, 813–845 10.1042/EBC20200067

2. Raetz, C. R., and Dowhan, W. (1990) Biosynthesis and function of phospholipids in Escherichia coli J Biol Chem 265, 1235–1238

3. Sohlenkamp, C., and Geiger, O. (2016) Bacterial membrane lipids: diversity in structures and pathways FEMS Microbiol Rev 40, 133–159 10.1093/femsre/fuv008

4. Donohue, T. J., Cain, B. D., and Kaplan, S. (1982) Alterations in the phospholipid composition of Rhodopseudomonas sphaeroides and other bacteria induced by Tris J Bacteriol 152, 595–606 10.1128/jb.152.2.595-606.1982

5. Sohlenkamp, C., Lopez-Lara, I. M., and Geiger, O. (2003) Biosynthesis of phosphatidylcholine in bacteria Prog Lipid Res 42, 115–162 10.1016/s0163-7827(02)00050-4

6. Geiger, O., Lopez-Lara, I. M., and Sohlenkamp, C. (2013) Phosphatidylcholine biosynthesis and function in bacteria Biochim Biophys Acta 1831, 503–513 10.1016/j.bbalip.2012.08.009

7. Kowalczyk, B., Chmiel, E., and Palusinska-Szysz, M. (2021) The Role of Lipids in Legionella-Host Interaction Int J Mol Sci 22, 10.3390/ijms22031487

8. Aktas, M., Wessel, M., Hacker, S., Klusener, S., Gleichenhagen, J., and Narberhaus, F. (2010) Phosphatidylcholine biosynthesis and its significance in bacteria interacting with eukaryotic cells Eur J Cell Biol 89, 888–894 10.1016/j.ejcb.2010.06.013

9. Wessel, M., Klusener, S., Godeke, J., Fritz, C., Hacker, S., and Narberhaus, F. (2006) Virulence of Agrobacterium tumefaciens requires phosphatidylcholine in the bacterial membrane Mol Microbiol 62, 906–915 10.1111/j.1365-2958.2006.05425.x

10. Minder, A. C., de Rudder, K. E., Narberhaus, F., Fischer, H. M., Hennecke, H., and Geiger, O. (2001) Phosphatidylcholine levels in Bradyrhizobium japonicum membranes are critical for an efficient symbiosis with the soybean host plant Mol Microbiol 39, 1186–1198

11. Aktas, M., Koster, S., Kizilirmak, S., Casanova, J. C., Betz, H., Fritz, C. et al. (2014) Enzymatic properties and substrate specificity of a bacterial phosphatidylcholine synthase FEBS J 281, 3523–3541 10.1111/febs.12877

12. de Rudder, K. E., Sohlenkamp, C., and Geiger, O. (1999) Plant-exuded choline is used for rhizobial membrane lipid biosynthesis by phosphatidylcholine synthase J Biol Chem 274, 20011–20016 10.1074/jbc.274.28.20011

13. Arondel, V., Benning, C., and Somerville, C. R. (1993) Isolation and functional expression in Escherichia coli of a gene encoding phosphatidylethanolamine methyltransferase (EC 2.1.1.17) from Rhodobacter sphaeroides J Biol Chem 268, 16002–16008

14. Klusener, S., Aktas, M., Thormann, K. M., Wessel, M., and Narberhaus, F. (2009) Expression and physiological relevance of Agrobacterium tumefaciens phosphatidylcholine biosynthesis genes J Bacteriol 191, 365–374 10.1128/JB.01183-08

15. Aktas, M., and Narberhaus, F. (2009) In vitro characterization of the enzyme properties of the phospholipid N-methyltransferase PmtA from Agrobacterium tumefaciens J Bacteriol 191, 2033–2041 10.1128/JB.01591-08

16. Aktas, M., Gleichenhagen, J., Stoll, R., and Narberhaus, F. (2011) S-adenosylmethionine-binding properties of a bacterial phospholipid N-methyltransferase J Bacteriol 193, 3473–3481 10.1128/JB.01539-10

17. Danne, L., Aktas, M., Gleichenhagen, J., Grund, N., Wagner, D., Schwalbe, H. et al. (2015) Membrane-binding mechanism of a bacterial phospholipid N-methyltransferase Mol Microbiol 95, 313–331 10.1111/mmi.12870

18. Danne, L., Aktas, M., Unger, A., Linke, W. A., Erdmann, R., and Narberhaus, F. (2017) Membrane Remodeling by a Bacterial Phospholipid-Methylating Enzyme mBio 8, 10.1128/mBio.02082-16

19. Danne, L., Aktas, M., Grund, N., Bentler, T., Erdmann, R., and Narberhaus, F. (2017) Dissection of membrane-binding and -remodeling regions in two classes of bacterial phospholipid N-methyltransferases Biochim Biophys Acta Biomembr 1859, 2279–2288 10.1016/j.bbamem.2017.09.013

20. Hacker, S., Sohlenkamp, C., Aktas, M., Geiger, O., and Narberhaus, F. (2008) Multiple phospholipid N-methyltransferases with distinct substrate specificities are encoded in Bradyrhizobium japonicum J Bacteriol 190, 571–580 10.1128/JB.01423-07

21. Kleetz, J., Vasilopoulos, G., Czolkoss, S., Aktas, M., and Narberhaus, F. (2021) Recombinant and endogenous ways to produce methylated phospholipids in Escherichia coli Appl Microbiol Biotechnol 105, 8837–8851 10.1007/s00253-021-11654-8

22. Kleetz, J., Welter, L., Mizza, A. S., Aktas, M., and Narberhaus, F. (2021) Phospholipid N-Methyltransferases Produce Various Methylated Phosphatidylethanolamine Derivatives in Thermophilic Bacteria Appl Environ Microbiol 87, e0110521 10.1128/AEM.01105-21

23. Salsabila, S. D., and Kim, J. (2024) Structural insights into phosphatidylethanolamine N-methyltransferase PmtA mediating bacterial phosphatidylcholine synthesis Sci Adv 10, eadr0122 10.1126/sciadv.adr0122

24. Martin, J. L., and McMillan, F. M. (2002) SAM (dependent) I AM: the S-adenosylmethionine-dependent methyltransferase fold Curr Opin Struct Biol 12, 783–793 10.1016/s0959-440x(02)00391-3

25. Jumper, J., Evans, R., Pritzel, A., Green, T., Figurnov, M., Ronneberger, O., et al. (2021) Highly accurate protein structure prediction with AlphaFold Nature 596, 583–589 10.1038/s41586-021-03819-2

26. Mirdita, M., Schutze, K., Moriwaki, Y., Heo, L., Ovchinnikov, S., and Steinegger, M. (2022) ColabFold: making protein folding accessible to all Nat Methods 19, 679–682 10.1038/s41592-022-01488-1

27. Holm, L., Laiho, A., Toronen, P., and Salgado, M. (2023) DALI shines a light on remote homologs: One hundred discoveries Protein Sci 32, e4519 10.1002/pro.4519

28. Fislage, M., Roovers, M., Tuszynska, I., Bujnicki, J. M., Droogmans, L., and Versees, W. (2012) Crystal structures of the tRNA:m2G6 methyltransferase Trm14/TrmN from two domains of life Nucleic Acids Res 40, 5149–5161 10.1093/nar/gks163

29. Kumar, A., Saigal, K., Malhotra, K., Sinha, K. M., and Taneja, B. (2011) Structural and functional characterization of Rv2966c protein reveals an RsmD-like methyltransferase from Mycobacterium tuberculosis and the role of its N-terminal domain in target recognition J Biol Chem 286, 19652–19661 10.1074/jbc.M110.200428

30. Lang, D. E., Morris, J. S., Rowley, M., Torres, M. A., Maksimovich, V. A., Facchini, P. J., et al. (2019) Structure-function studies of tetrahydroprotoberberine N-methyltransferase reveal the molecular basis of stereoselective substrate recognition J Biol Chem 294, 14482–14498 10.1074/jbc.RA119.009214

31. Lee, S. G., Kim, Y., Alpert, T. D., Nagata, A., and Jez, J. M. (2012) Structure and reaction mechanism of phosphoethanolamine methyltransferase from the malaria parasite Plasmodium falciparum: an antiparasitic drug target J Biol Chem 287, 1426–1434 10.1074/jbc.M111.315267

32. Morana, A., Stiuso, P., Colonna, G., Lamberti, M., Carteni, M., and De Rosa, M. (2002) Stabilization of S-adenosyl-L-methionine promoted by trehalose Biochim Biophys Acta 1573, 105–108 10.1016/s0304-4165(02)00333-1

33. Trott, O., and Olson, A. J. (2010) AutoDock Vina: improving the speed and accuracy of docking with a new scoring function, efficient optimization, and multithreading J Comput Chem 31, 455–461 10.1002/jcc.21334

34. Lee, S. G., and Jez, J. M. (2013) Evolution of structure and mechanistic divergence in di-domain methyltransferases from nematode phosphocholine biosynthesis Structure 21, 1778–1787 10.1016/j.str.2013.07.023

35. Lee, S. G., and Jez, J. M. (2017) Conformational changes in the di-domain structure of Arabidopsis phosphoethanolamine methyltransferase leads to active-site formation J Biol Chem 292, 21690–21702 10.1074/jbc.RA117.000106

36. de Rudder, K. E., Thomas-Oates, J. E., and Geiger, O. (1997) Rhizobium meliloti mutants deficient in phospholipid N-methyltransferase still contain phosphatidylcholine J Bacteriol 179, 6921–6928 10.1128/jb.179.22.6921-6928.1997

37. Moser, R., Aktas, M., and Narberhaus, F. (2014) Phosphatidylcholine biosynthesis in Xanthomonas campestris via a yeast-like acylation pathway Mol Microbiol 91, 736–750 10.1111/mmi.12492

38. Kodaki, T., and Yamashita, S. (1987) Yeast phosphatidylethanolamine methylation pathway. Cloning and characterization of two distinct methyltransferase genes J Biol Chem 262, 15428–15435

39. McGraw, P., and Henry, S. A. (1989) Mutations in the Saccharomyces cerevisiae opi3 gene: effects on phospholipid methylation, growth and cross-pathway regulation of inositol synthesis Genetics 122, 317–330 10.1093/genetics/122.2.317

40. Ridgway, N. D., and Vance, D. E. (1988) Kinetic mechanism of phosphatidylethanolamine N-methyltransferase J Biol Chem 263, 16864–16871

41. Hirata, K., Yamashita, K., Ueno, G., Kawano, Y., Hasegawa, K., Kumasaka, T. et al. (2019) ZOO: an automatic data-collection system for high-throughput structure analysis in protein microcrystallography Acta Crystallogr D Struct Biol 75, 138–150 10.1107/S2059798318017795

42. Yamashita, K., Hirata, K., and Yamamoto, M. (2018) KAMO: towards automated data processing for microcrystals Acta Crystallogr D Struct Biol 74, 441–449 10.1107/S2059798318004576

43. Kabsch, W. (2010) Xds Acta Crystallogr D Biol Crystallogr 66, 125–132 10.1107/S0907444909047337

44. McCoy, A. J., Grosse-Kunstleve, R. W., Adams, P. D., Winn, M. D., Storoni, L. C., and Read, R. J. (2007) Phaser crystallographic software J Appl Crystallogr 40, 658–674 10.1107/S0021889807021206

45. Emsley, P., Lohkamp, B., Scott, W. G., and Cowtan, K. (2010) Features and development of Coot Acta Crystallogr D Biol Crystallogr 66, 486–501 10.1107/S0907444910007493

46. Afonine, P. V., Grosse-Kunstleve, R. W., Echols, N., Headd, J. J., Moriarty, N. W., Mustyakimov, M. et al. (2012) Towards automated crystallographic structure refinement with phenix.refine Acta Crystallogr D Biol Crystallogr 68, 352–367 10.1107/S0907444912001308

47. Williams, C. J., Headd, J. J., Moriarty, N. W., Prisant, M. G., Videau, L. L., Deis, L. N. et al. (2018) MolProbity: More and better reference data for improved all-atom structure validation Protein Sci 27, 293–315 10.1002/pro.3330

48. Delaglio, F., Grzesiek, S., Vuister, G. W., Zhu, G., Pfeifer, J., and Bax, A. (1995) NMRPipe: a multidimensional spectral processing system based on UNIX pipes J Biomol NMR 6, 277–293 10.1007/BF00197809

49. Kazimierczuk, K., and Orekhov, V. Y. (2011) Accelerated NMR spectroscopy by using compressed sensing Angew Chem Int Ed Engl 50, 5556–5559 10.1002/anie.201100370

50. Watanabe, Y., Watanabe, Y., and Watanabe, S. (2020) Structural Basis for Phosphatidylethanolamine Biosynthesis by Bacterial Phosphatidylserine Decarboxylase Structure 28, 799–809 e795 10.1016/j.str.2020.04.006

51. Sievers, F., Wilm, A., Dineen, D., Gibson, T. J., Karplus, K., Li, W., et al. (2011) Fast, scalable generation of high-quality protein multiple sequence alignments using Clustal Omega Mol Syst Biol 7, 539 10.1038/msb.2011.75

52. Robert, X., and Gouet, P. (2014) Deciphering key features in protein structures with the new ENDscript server Nucleic Acids Res 42, W320–324 10.1093/nar/gku316

