## Supporting Information for "Structural basis for phosphatidylcholine synthesis by bacterial phospholipid *N*-methyltransferases"

From [1]Faculty of Science, Yamagata University, 1-4-12 Kojirakawa-machi, Yamagata 990-8560, Japan; [2]Graduate School of Life Science, Hokkaido University, Kita 10, Nishi 8, Kita-ku, Sapporo, Hokkaido 060-0808, Japan; [3]Faculty of Agriculture, Ehime University, 3-5-7 Tarumi, Matsuyama, Ehime, 790-8566, Japan; [4]Department of Bioscience, Graduate School of Agriculture, Ehime University, 3-5-7 Tarumi, Matsuyama, Ehime, 790-8566, Japan; [5]Center for Marine Environmental Studies (CMES), Ehime University, 2-5 Bunkyo-cho, Matsuyama, Ehime 790-8577, Japan.

*Corresponding author: Yasunori Watanabe; Faculty of Science, Yamagata University, 1-4-12 Kojirakawa-machi, Yamagata 990-8560, Japan;

**Materials included:**

Figure S1

Figure S2

Figure S3

Figure S4

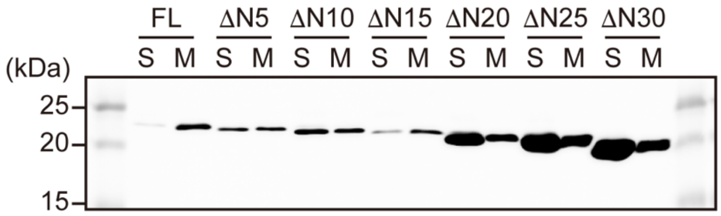


**Figure S1. Membrane-binding activity of the N-terminal region lacking AtPmtA mutants**
Soluble (S) and membrane (M) fractions were separated from *E. coli* cells expressing full-length AtPmtA (FL) or the N-terminal truncated mutants of AtPmtA (ΔN5, ΔN10, ΔN15, ΔN20, ΔN25, and ΔN30) and subjected to SDS-PAGE, followed by immunoblotting with anti-6x histidine antibody.

A

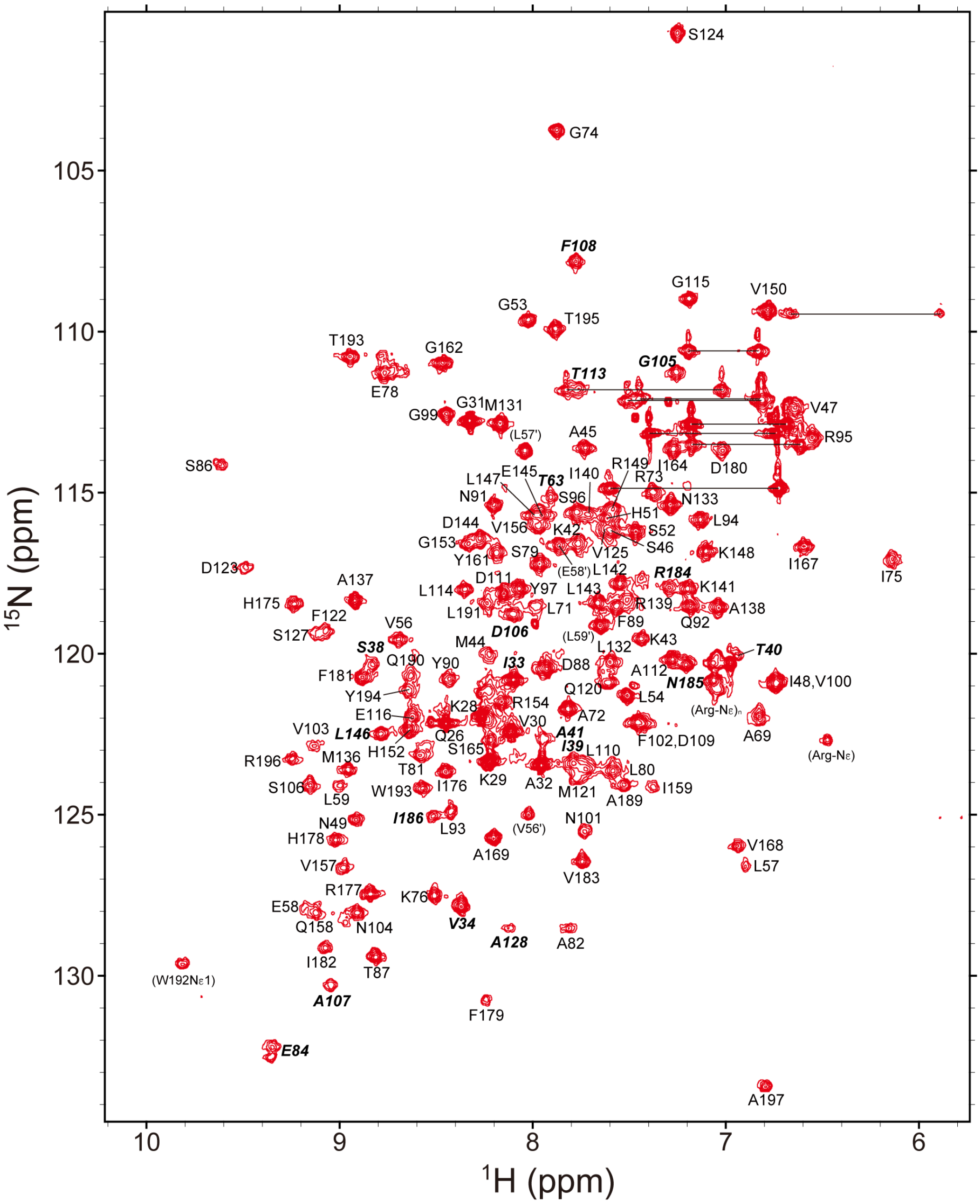


```
 26 QPKKVGAIVPTSSITAKKMASVINPHSGLPVLELGPGTGVITKAILARGIKPESL  80
 81 TAIEYSTDFYNQLLRSYPGVNFVNGDAFDLDATLGEHKGQMFDSVISAVPMLNFP  135
136 MAARIKLLDELLKRVPHGRPVVQISYGPISPIVAQPHLYHIRHFDFIVRNIPPAQ  190
191 LWTYTRA 197
```

B

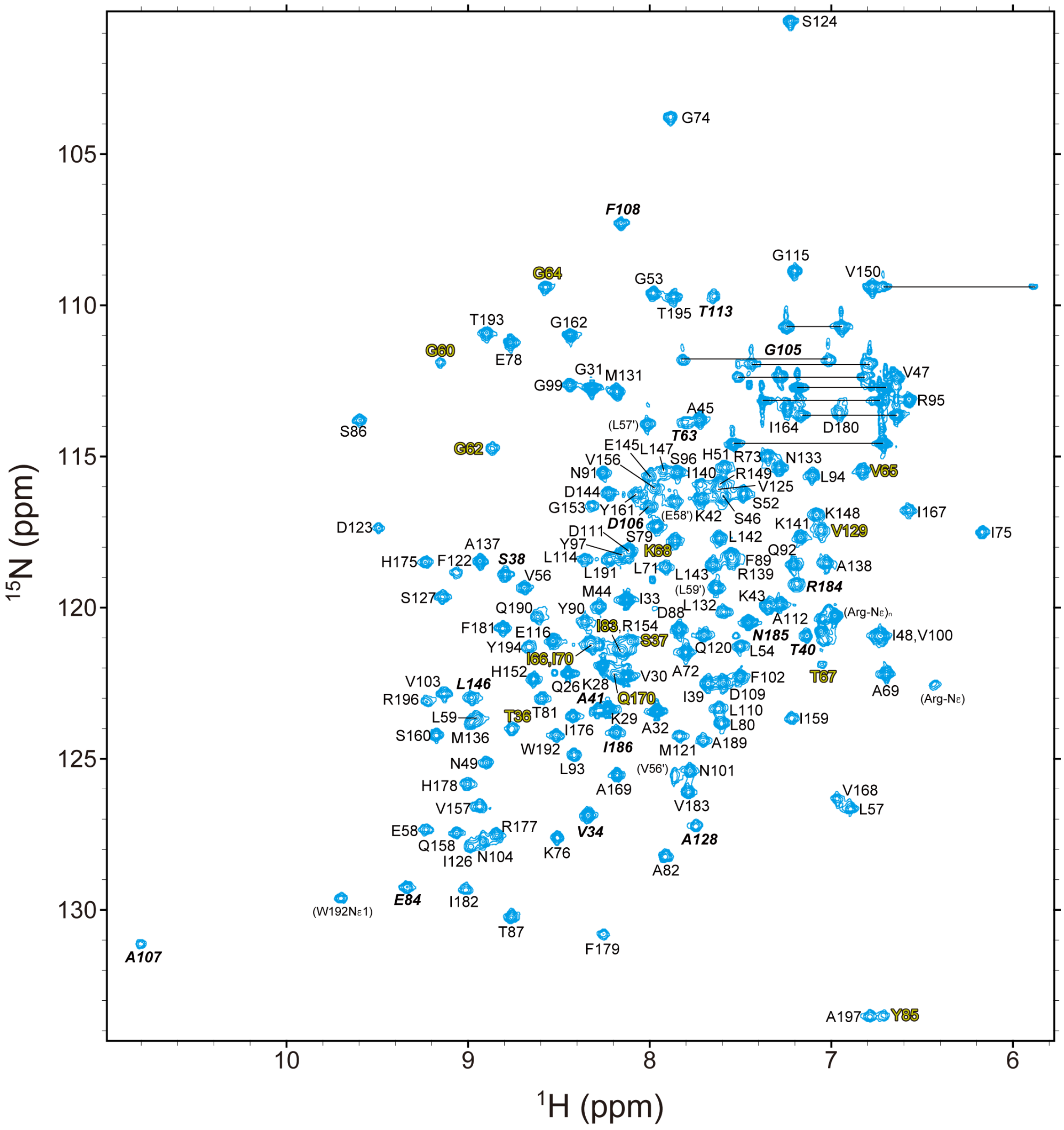


```
 26 QPKKVGAIVPTSSITAKKMASVINPHSGLPVLELGPGTGVITKAILARGIKPESL  80
 81 TAIEYSTDFYNQLLRSYPGVNFVNGDAFDLDATLGEHKGQMFDSVISAVPMLNFP  135
136 MAARIKLLDELLKRVPHGRPVVQISYGPISPIVAQPHLYHIRHFDFIVRNIPPAQ  190
191 LWTYTRA 197
```

**Figure S2. Assigned $^{1}$H-$^{15}$N HSQC spectra of apo and SAH bound AtPmtAΔN25**

(A-B) [$^{1}$H-$^{15}$N] HSQC spectra with resonance assignments of 300 μM AtPmtAΔN25 in the absence (A) and presence of 3 mM SAH (B). The residues with significant chemical shift changes upon binding to SAH are labeled in italic. The residues whose signals appeared upon binding to SAH are labeled in yellow. Minor form assignments are indicated by the prime symbol ('). Amino group resonances of Asn and Gln side chains are connected by horizontal lines. Amino acid sequence of AtPmtAΔN25, with assigned residues in black and unassigned residues in gray, is shown below each spectrum.

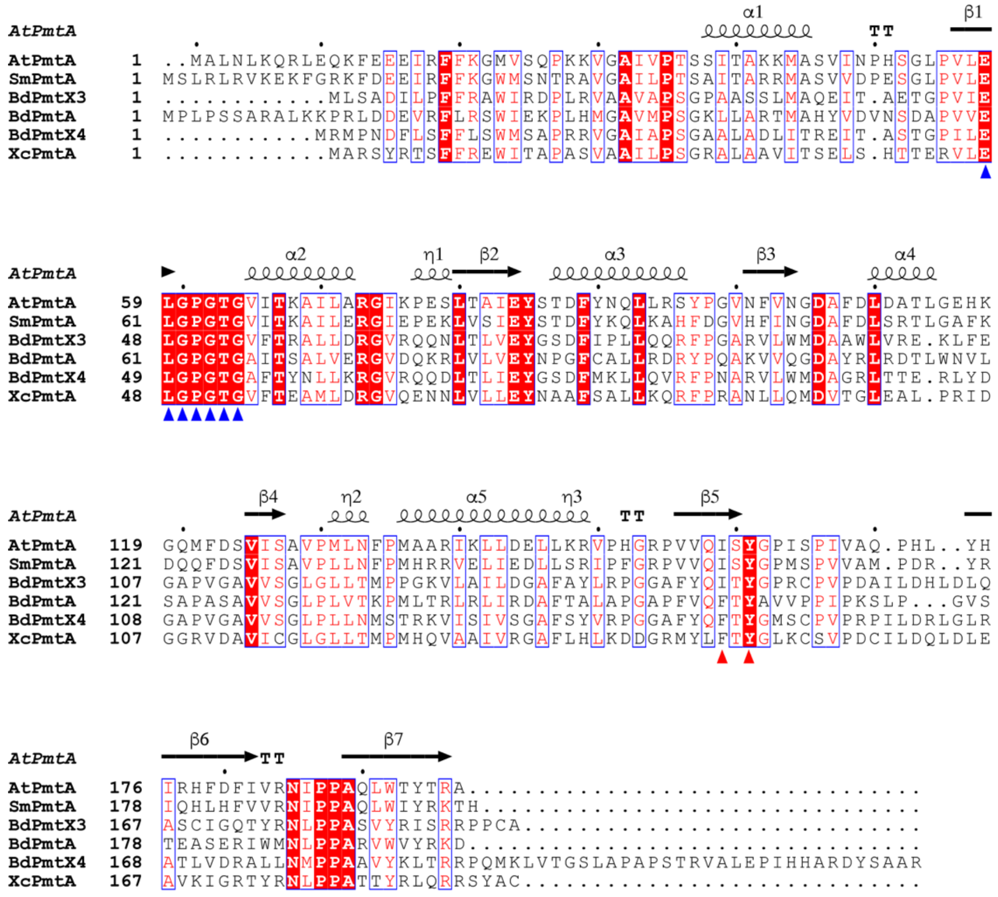

**Figure S3. Sequence alignment of S-type Pmt homologs**

Multiple-sequence alignment of AtPmtA with S-type Pmt homologs from *Sinorhizobium meliloti* (SmPmtA), *Bradyrhizobium diazoefficiens* (BdPmtA, BdPmtX3, and BdPmtX4), and *Xanthomonas campestris* (XcPmtA). Identical residues are shaded red, similar residues are colored red, and similar residues across groups are framed in blue. The secondary structure of AtPmtA is shown above the sequences. The conserved SAM-binding motifs are indicated with blue arrowheads. Ile159 and Tyr161 in AtPmtA are indicated with red arrowheads.

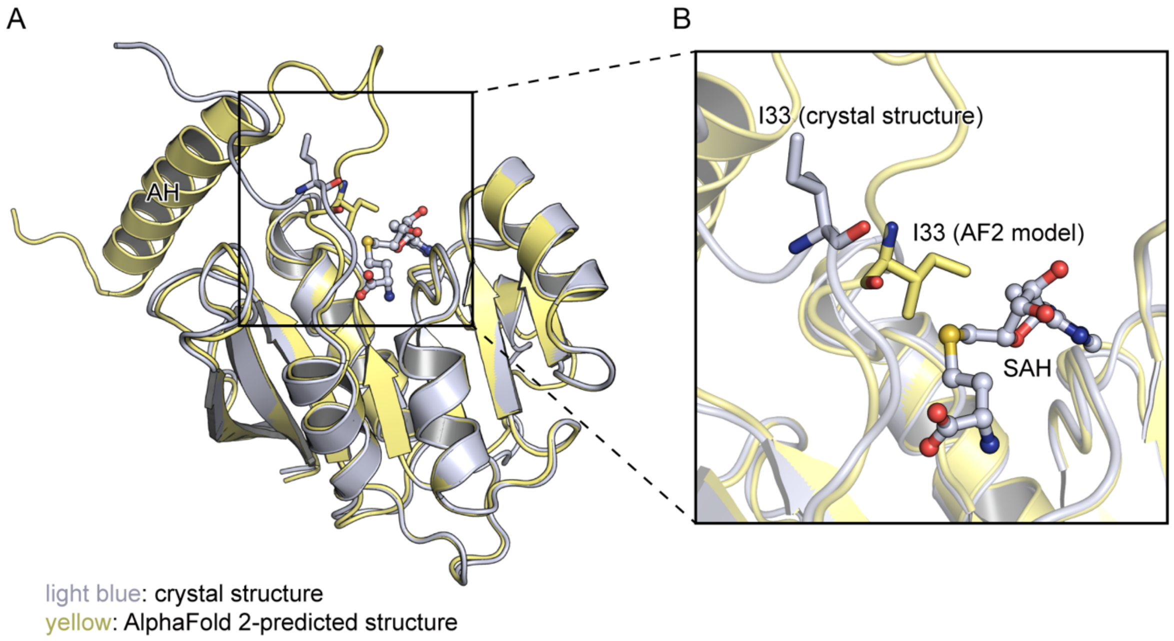


**Figure S4. Comparison of AlphaFold 2-predicted and crystal structures of AtPmtA**
(A) Superposition of the AlphaFold 2-predicted structure of AtPmtA (yellow) and the crystal structure of AtPmtAΔN25 (light blue). The N-terminal amphipathic helix of the AlphaFold 2-predicted structure of AtPmtA is labeled as AH. (B) The magnified view around Ile33 of AtPmtA. Ile33 is shown in stick form, whereas SAH is shown in stick and ball form.
